# Development and validation of a type 2 diabetes machine learning classification model for EHR-based diagnostics and clinical decision support

**DOI:** 10.1101/2022.10.08.511400

**Authors:** Victor Glanz, Vladimir Dudenkov, Alexey Velikorodny

**Author notes:** equal contributors (Vladimir Dudenkov, Alexey Velikorodny).

## Abstract

**Background:** Undiagnosed type 2 diabetes continues to represent a significant challenge for all national healthcare systems. Although diagnostic criteria and laboratory screening procedures are well-established, clinical tests have limitations, and in many cases, diagnosis confirmation and more accurate interpretation of the test results are required. Machine learning methods, when applied to clinical outcome risk prediction, demonstrate great effectiveness, as they recognize specific patterns in data dynamics and thus can be used for the identification of at-risk cases where diabetes and complications can be delayed or even prevented. The aim of this study was to develop a type 2 diabetes machine learning model capable of efficient early identification of diabetes presence based on the results of common laboratory tests.

**Methods:** Real-world medical data from electronic medical records were subjected to a multistage processing, including feature selection, missing values imputation. The machine learning algorithms adopted in this study were XGBoost, multilayer perceptron, ridge classifier, ridge classifier with polynomial features, bootstrap aggregating, dynamic ensemble selection, stacked generalization. An external dataset was analyzed via the same workflow to validate the initial results. The study was designed in accordance with the TRIPOD statement.

**Results:** We have developed a machine learning classification model for type 2 diabetes that possesses several important advantages over conventional clinical methods (specifically, FINDRISC, ADA risk score). Performance metrics for the diabetes diagnostic model were 0.96 AUC, 92% specificity, and 89% sensitivity (mean values).

**Conclusions:** The study results potentially have major clinical implication and provide a contribution to the field of conventional diabetes risk assessment tools. Being specifically trained on real-world laboratory data and based on satisfactory external validation results, the present diagnostic type 2 diabetes model demonstrates high generalizability and can serve as a medical decision support and health monitoring tool.

## Background

Diabetes remains one of the most rapidly developing health emergencies around the world despite continuous scientific progress uncovering the molecular basis of its pathogenesis. Globally, more than 537 million people are affected by diabetes, and the situation is expected to get even worse, as 46% more cases are predicted to emerge by 2045, according to the International Diabetes Federation (1). Being a complex systemic disorder, diabetes can be categorized into four general types, as proposed by the American Diabetes Association (ADA): 1) type 1 diabetes (T1D), absolute insulin deficiency resulting from autoimmune destruction of β-cells; 2) type 2 diabetes (T2D), loss of insulin secretion by β-cells and tissue sensitivity to insulin resulting from inflammation, metabolic stress and genetic predisposition to certain abnormalities of metabolism; 3) gestational diabetes (GA), pregnancy-specific type, not evident before gestation; 4) other specific types (2). Hyperglycemia is a clinical manifestation of various underlying mechanisms of diabetes development. Diagnostic procedures consist of laboratory measurements of plasma glucose (fasting or oral glucose tolerance test) and glycated hemoglobin. Current diagnostic cut-off values, established by ADA, are above 126 mg/dL (7.0 mmol/L) for fasting glucose and ≥ 6.5% for glycated hemoglobin.

Missed undiagnosed cases of diabetes and inability to efficiently detect high-risk individuals early are the main healthcare issues, that scientific research is yet to resolve. To address this problem, the use of machine learning and deep learning methods that created opportunities to improve medical data processing, which is confirmed by numerous studies (3,4). Such methods as support vector machine (SVM), random forest (RF), and extreme gradient boosting (XGBoost) have successfully been used to predict the outcomes for hypertension (5). Neural networks and deep learning techniques can be applied for electronic health records (EHR) analysis (6). The cooperative application of deep learning models and bootstrap aggregation was able to improve the quality of information extraction systems from cancer pathology reports (7). In managing diabetes, machine learning (ML) methods can facilitate both diagnosis and prognosis. Comprehensive literature reviews by Kodama et al. and Silva et al. provide an outlook of the most clinically relevant ML diabetes prediction models that were evaluated in terms of performance metrics (8,9). Prognostic only ML diabetes models had a mean area under the curve (AUC) value of 0.88, while pooled prognostic and diagnostic low-bias-algorithms showed 0.830 for c-index. In contrast, several traditional diabetes risk estimation methods (FINDRISC, AUSDRISC, etc.) showed a mean AUC of 0.79 (8). To improve diabetes diagnosis procedure, a diagnostic ML-system was proposed by Calisir and Dogantekin (10). Efficient diabetes classification was achieved via the stages of feature selection (by using the Linear Discriminant Analysis method), classification (by using Morlet Wavelet Support Vector Machine classifier), and diagnostic performance evaluation. Although high values of sensitivity and specificity were obtained (0.833 and 0.937, respectively), the system was not validated on real-world data (the Pima Indians Diabetes Database was used for the study). Zheng et al. proposed a semi-automated framework based on machine learning and feature engineering (11). Lab features were supplemented with information on self-reported diabetic symptoms and complications. The framework included six machine-learning models: k-Nearest-Neighbors, Naïve Bayes, Decision Tree, Random Forest, Support Vector Machine, and Logistic Regression. EHR data generated by ten local EHR systems were used for the training and validation stages of the study. The authors reported an average AUC of 0.98 during 4-fold cross-validation. Han et al. used support vector machines to screen for diabetes and added an ensemble learning module, which turned SVM decisions into an explicit set of rules (12). The considered feature set was limited to nine measurements: glycated hemoglobin A1c, triglycerides, uric acid, high-density lipoprotein, age, diastolic blood pressure, total cholesterol, waist circumference, and weight. The data from the China Health and Nutrition Survey was used as the data source. Rule sets generated by the ensemble learning method demonstrated weighted average precision of 0.942 and an average recall of 0.939. Nguyen et al. presented a deep learning model that combined wide deep classifiers and an ensemble model to improve diabetes prediction (13). A wide range of features was considered, including diagnoses, medications, and laboratory tests. The authors used a publicly available American EHR dataset released by Practice Fusion in 2012 for a data science competition. During 10-fold cross-validation the following performance metrics were achieved: AUC 0.841, sensitivity 0.311, and specificity 0.968.

Type 2 diabetes was the focus of the present work due to several reasons. First, T2D affects significantly more individuals than any other type and represents 90%-95% of all cases. Worth noting that T2D is no longer considered as a ‘disease of the adults’, nor does T1D occur specifically at an early age. Second, the rate of undiagnosed T2D cases is alarmingly high due to the increase in plasma glucose levels being gradual, often too slow for a person to notice any symptoms. Finally, although T2D screening laboratory tests and risk prediction algorithms are established and available in the clinic, the proper time interval between screenings is not defined. Current ADA guidelines recommend a minimum 3-year interval, but it is admitted that the timing can and should be personalized (2). Personalized screening intervals based on diabetes risk can be critical for timely intervention aimed at delaying or preventing the onset of T2D.

Overall, while previous studies established a foundation to build efficient diabetes models, several issues that substantially reduce their performance remain unresolved: 1) use of subjective information, such as patient-reported symptoms and medical history instead of precise laboratory data; 2) lack of personalized data-based real-time feature selection - most of the existing models use limited sets of predicting features, chosen primarily from literature; 3) diabetes and other disease-specific models are usually trained on clinical study data with little to no missing parameters. However, missing data are omnipresent in a real-world setting. Therefore, patients with incomplete data are usually excluded from modeling studies; 4) common lack of external validation limits translation of study results into practice. The current study addresses these issues by combining real-world patient-generated laboratory data from EHR, automatic feature selection approach, widely used imputation methods, and sophisticated machine learning algorithms in a prediction model that can serve as a medical decision support and health monitoring tool.

## Methods

### Data source and data selection

The source of data for the study were EHRs containing ultrasound markers, laboratory measurement results and information about clinical outcomes (ICD-10 codes determined by medical specialists). We used records from 04.09.2018 to 01.01.2022 extracted from a private anonymized dataset that included 157615 patients. After primary filtration, the following patients were excluded: T1D and other non-T2D types of diabetes, pregnant women, and patients under 18 years of age. The remaining data underwent the outlier detection and feature selection procedures. Only the information regarding patients with significant (relevant) features was obtained. The following selection strategy was applied at the next stage: measurement results were gathered 12 months before the diagnosis for each patient in the ‘diabetes’ cohort and 12 months before the examination by an endocrinologist for non-diabetics. Table rows that did not contain information about fasting plasma glucose level were excluded. The described data filtration process is shown in Fig. 1.

**Figure 1.**
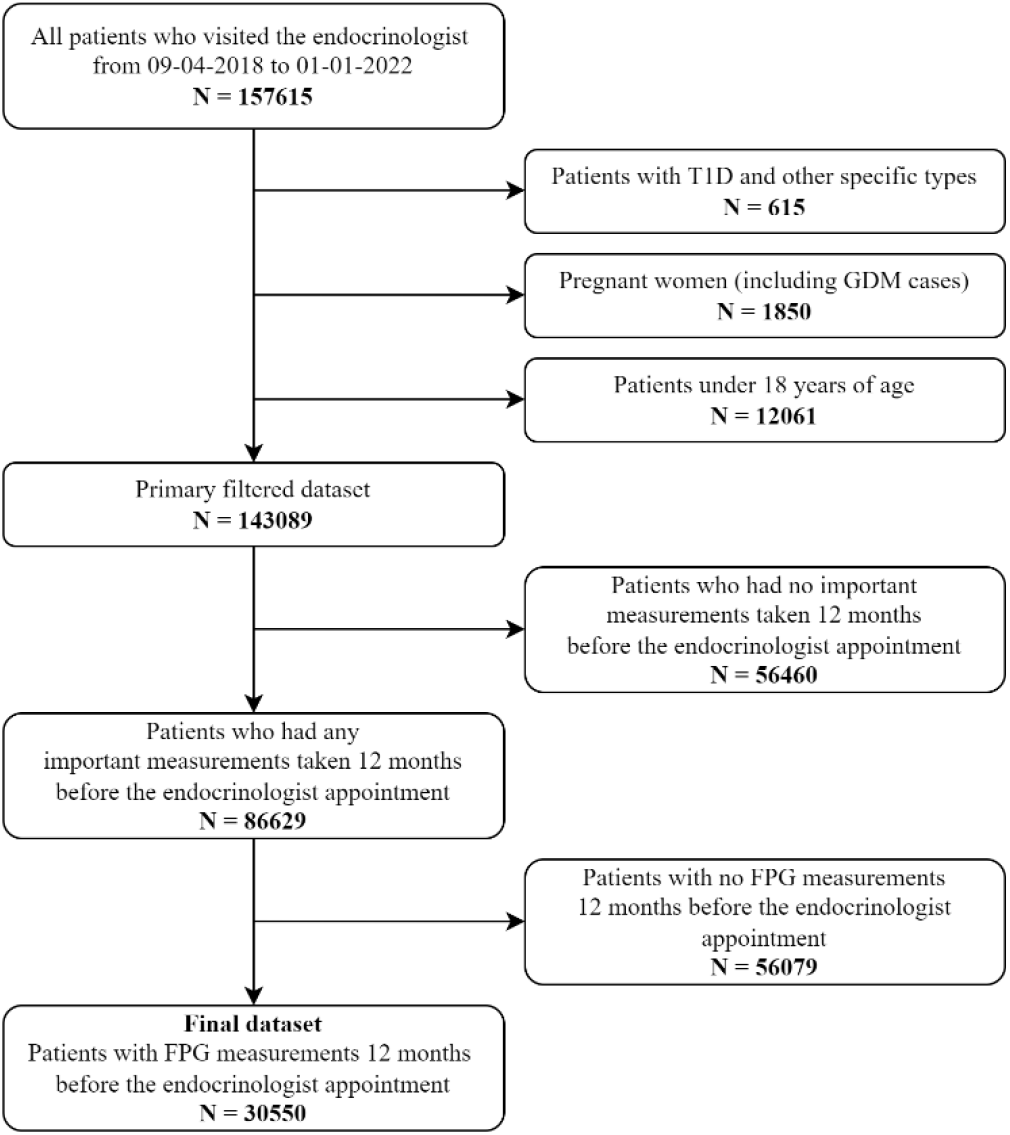
Data selection chart for the main dataset

### Model development workflow

EHR data went through the sequential stages of collection, feature selection, and outlier detection. The resulting dataset was split into test and training subsets. We used a stratified split of 70% and 30% for the training and testing datasets, respectively. With the stratified split strategy, the ratio of diabetics to non-diabetic individuals in splits remains the same as in the full population. Since the splitting was random, it was repeated three times to ensure the reliability of the results. Because of missing values in the data, it was necessary to apply data imputation methods (14). Both multivariate and univariate approaches to missing-values imputation were considered in this work. The data imputation step was combined with the hyperparameter optimization procedure. A specific imputation method was individually selected for each machine learning model. The standard procedure of removing mean values and normalizing to unit variance was then applied.

The following algorithms were selected, based on the current trends in machine learning (15,16):

- XGBoost
- Multilayer perceptron
- Ridge classifier
- Ridge classifier with polynomial features
- Bootstrap aggregating
- Dynamic ensemble selection.

Additionally, a stacked generalization was employed as a combination of several machine learning models, with logistic regression as a meta-estimator. Final modeling results were evaluated and interpreted both by biomedical experts and statistical methods. The research workflow shown in Fig. 2 was designed in accordance with the TRIPOD (Transparent Reporting of a multivariable prediction model for Individual Prognosis or Diagnosis) statement (17).

**Figure 2.**
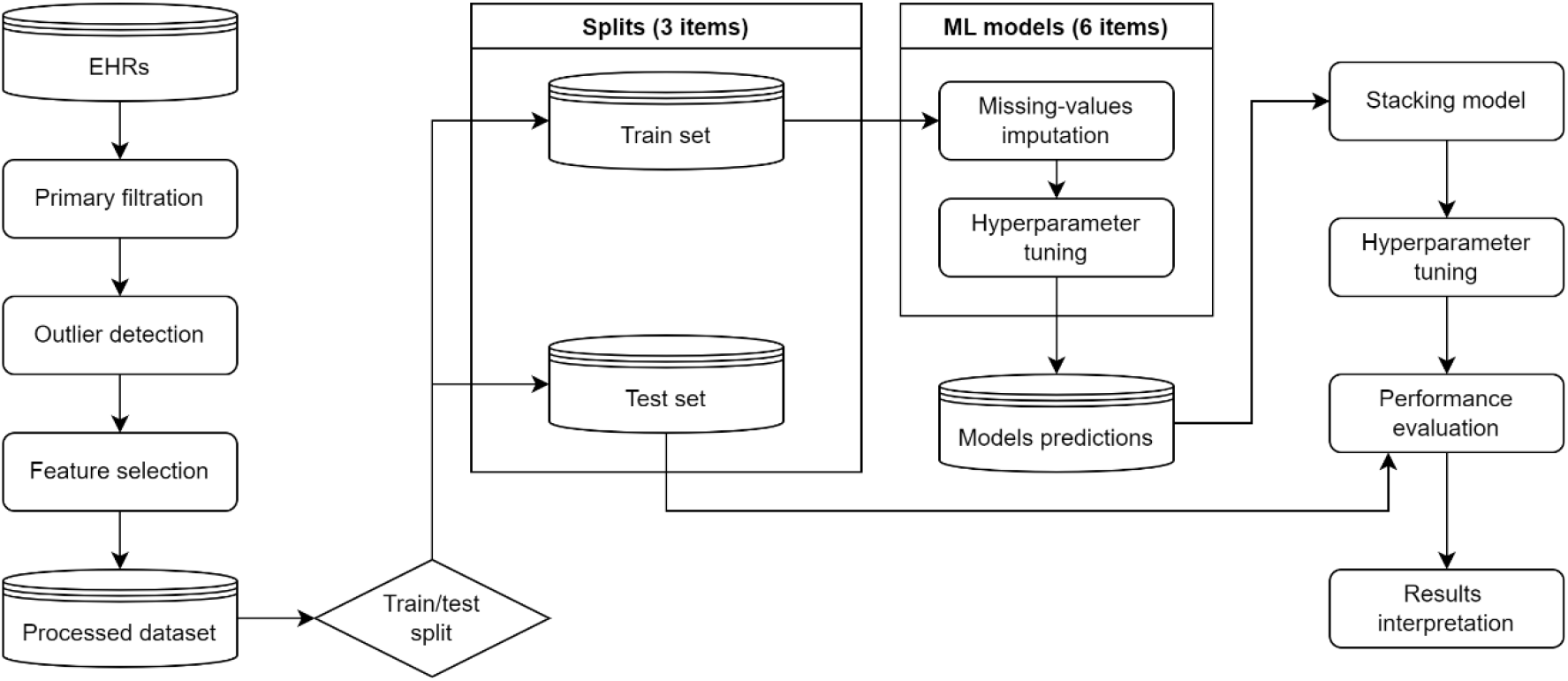
Workflow chart of the diagnostic model development

#### Outlier detection

Prior experience with clinical biomarker data had shown that automatic methods did not provide sufficient quality for outlier detection. Thus, ‘cleaning’ the data of outliers was carried out using the intervals of acceptable values specified by laboratory manuals and results of literature analysis.

#### Feature selection

Feature selection is the process of dimensionality reduction of the feature space that helps to improve computation time and to better understand the complex data (18,19). In addition, a possible reduction of features number is necessary for medical application of the model. Feature importance is the central concept of feature selection and refers to a score assigned to all the input features. A higher score means that the feature will have a larger effect on the ML model used to predict a target variable. The feature selection methods described hereafter have been proved to be efficient in the following studies (20–22).

##### Mutual information

Mutual information is a non-negative measure from the field of information theory. It is calculated between two variables and measures the amount of information one can obtain from a random variable from another. The formal definition for mutual information between two random variables *X* and *Y* is as follows:

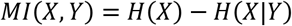

where (*MI*(*X*, *Y*)) is the mutual information for *X* and *Y*, *H*(*X*) is the entropy for *X* and *H*(*X*|*Y*) is the conditional entropy for *X* given *Y*.

Mutual information is straightforward when discrete variables are considered. However, it can be adapted to a continuous input with a discrete output. Nonparametric methods are used in this case, based on the entropy estimate for k-nearest neighboring distances (23,24).

##### The Gini index

A well-known measure of inequality, the Gini index, can be used for the feature selection task (25). It measures the ‘purity’ of features in relation to a class. Gini index defines the discrimination level of a feature to distinguish between possible classes. The Gini index for a feature *F* is defined as:

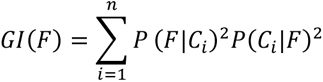

where *P*(*F*|*C_i_*) is the feature *F* probability given class *C_i_*, *P*(*C_i_*|*F*) is the class *C_i_* probability given the feature *F*. In our case, respectively, n = 2. Gini index measures the ‘purity’ when using a chosen feature, and assumes values in the range [0,1]. Therefore, the value 1 – *GI* was used as a feature importance measure.

#### ANOVA

Analysis of variance (ANOVA) is a collection of statistical methods to analyze the differences between the means of groups (26). The ANOVA F-value is:

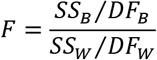

where *SS_B_* is a sum of squares between the groups, *DF_B_* is a degree of freedom for a mean square between the groups, *SS_W_* is a sum of squares within the groups, *DF_W_* is a degree of freedom for a mean square within the groups. High F-values correspond to the best discriminative capacity of the feature.

##### Tree-based method

Extremely randomized trees were employed as an intrinsic tree-based method (27). It is an ensemble learning technique that aggregates the outcomes of many decision trees to generate the final prediction. In case of tree-based method, the importance of a feature is computed as the normalized total reduction of the criterion brought about by that feature (impurity-based importance). Because of the stochastic nature of the algorithm, feature importance values might vary between calculations. Therefore, the final value was an average from ten runs.

##### Combining results

To generalize the results of all methods, the total importance of the features was computed. The selected methods were applied to the feature space, resulting in a M×N matrix, where M is the number of features, N is the number of methods. The values in each row were normalized and added up. The final values represent the generalized feature importance.

#### Data imputation and hyperparameter optimization

Electronic health records (EHRs) often include a high proportion of missing data, which can create difficulties in the implementation of machine learning models. Most algorithms require that their inputs have no missing values. A straightforward way to work with incomplete EHRs is to discard entire records containing missing values. However, this approach dictates a considerable risk of losing the significant amount of information necessary for efficient model performance. Here, this approach is unacceptable since the developed model is aimed at real clinical practices, where one must always deal with missing data. Thus, the imputation methods for clinical data were employed (28,29). To achieve the best result, the data imputation step was combined with the hyperparameter tuning. It was assumed that each model can have its optimal data imputation algorithm. Therefore, the data imputation method and its parameters were considered as additional hyperparameters. It should be noted that XGBoost model, for instance, does not require data imputation procedures. Data imputation types are listed below. Both univariate and multivariate imputation methods were employed.

- K-nearest neighbor (KNN). A missing sample is imputed by finding the closest samples in the training set and averaging these nearby samples to fill in the value. The configuration of KNN imputation involves selecting the distance measure and the number of neighbors for each prediction.
- Multiple Imputation by Chained Equations (MICE). MICE imputes missing values through an iterative series of predictive models (30). In each iteration, each specified feature in the dataset is imputed using other features. Iterations must be conducted until convergence is achieved.
- Univariate imputation using the feature statistics. This method assumes that the missing values are replaced with a specific statistical measure corresponding to the entire feature column (median, mode, or mean value). The choice of a particular statistic is a configurable parameter of this method.

The listed imputation methods are also often used in medical and bioinformatics research (31–33). Additionally, all methods were supplemented with a missing-indicator (34). Each dimension was marked with a Boolean flag indicating whether its value is missing or not.

A state-of-the-art Optuna framework was used for hyperparameter optimization (35). Tree-structured Parzen Estimator (TPE) was chosen as a sampling method (36). TPE is a single-objective Bayesian algorithm that uses tree-structured adaptive Parzen estimators as a surrogate (37).

#### Machine learning algorithms

##### XGBoost

Extreme Gradient Boosting (XGBoost) is a state-of-the-art algorithm designed to be parallelizable, computationally efficient, and resistant to overfitting. It is based on the idea of gradient boosting (38). Gradient boosting combines the predictions of weak estimators by dinting additive training strategies. The classification and regression trees (CART) are used as weak estimators in XGBoost. The output of each CART is combined using the following additive function:

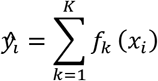

where *K* is the number of trees, *f_k_* is a function in the functional space of all possible CARTs. XGBoost learning is an iterative process. At every iteration, a new CART tries to rectify the weakness in the prediction of a previous one. 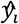 for iteration t is given as:

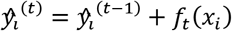

where 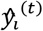 is the prediction for iteration *t*, *f_t_*(*x_i_*) is an additive function for iteration *t*. XGBoost introduces the idea of regularization to reduce the model complexity. The objective with regularization is:

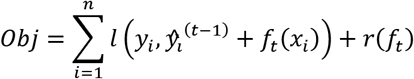

where *l* is the loss function, *r* is the regularization term. The regularization term is defined as:

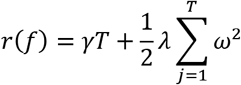

where *γ* and *λ* are the regularization degree control parameters, *T* is the number of leaves, *ω* is the score on each leaf. The regularization and tree-specific parameters, the number of boosting iterations, and sampling restrictions were selected by the hyperparameters tuning procedure.

##### Multilayer perceptron

A multilayer perceptron (MLP) is a fully connected artificial neural network (39). MLP consists of layers of neurons: one input layer, one or more hidden layers, and one output layer. Each neuron receives inputs from other neurons of the previous layer and provides outputs to the neurons of the following layer. The presence of at least one hidden layer allows MLP to approximate any continuous function. The connection strength between neurons is controlled by weights. The output of a given neuron is a linear weighted sum of inputs from the previous layer, which is passed through the activation function:

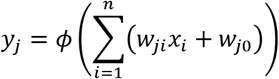

where *y_j_* is an output of j-th neuron, *x_i_* is an input from the i-th neuron of the previous layer, *W_ji_* is weight, *ϕ* is the activation function.

Weights are obtained via the training process by optimizing a predefined loss function. We used the binary cross entropy loss function:

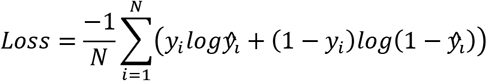

where 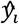 is the i-th value in the model output, *y_i_* is the corresponding target value, *N* is the number of values in the model output. Here, *N = 1*.

The dropout can be used as a neural network regularization method (40). It improves the generalization performance by repeating the procedure of temporarily removing neurons. The choice of neurons to be dropped is random. The strength of dropout can be regulated by the dropout rate, i.e., by the probability that the neuron will be removed.

The dropout rate, an activation function, the number of hidden layers, and the number of neurons in layers were the main hyperparameters for tuning the present MLP model.

##### Ridge classifier

Ridge regression improves the classic linear regression model by imposing L2 regularization element (41). The ridge coefficients minimize penalized residual sum of squares:

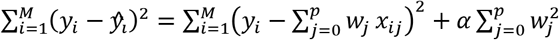

where *α* is a regularization strength. Higher values specify stronger regularization. We identified regularization strength as one of the most important for the hyperparameter optimization stage. The ridge classifier, based on the ridge regression method, can be applied to the binary classification problem. It converts the target variable (“diabetic” and “non-diabetic” in this case) into [-1, 1] and applies the regression method. The highest value in prediction is accepted as a target class.

##### Ridge classifier with polynomial features

The polynomial extension of the ridge classifier was considered as a separate model. We expanded the feature space by polynomial combinations of several base features with degrees less than or equal to the specified limit. Thereby, the specified degree and the set of features to expand functioned as parameters for the hyperparameter optimization. The described polynomial extension of the ridge classifier may be considered a type of feature engineering. This extension allows to identify nonlinear patterns (42).

##### Bootstrap aggregating

Bootstrap aggregation (bagging) is an ensemble learning technique that uses bootstrap sampling (43). The bagging fits base estimators on different bootstrap samples of the training dataset and combines them using the average of the predictions, or a different statistic. Bagging is a classic machine learning model used frequently to improve the accuracy of arbitrary base estimators (44,45). The type of base estimators and their number were significant parameters of the algorithm, hence their inclusion in the hyperparameter optimization stage. Bootstrap parameters, such as the sample limit and the feature limit for the splitting, were also employed.

##### Dynamic ensemble selection

Dynamic ensemble selection strategies refer to the ensemble learning techniques that dynamically choose a subset of models to make a prediction based on a specific input (46). The subset is defined according to the model competence, estimated over certain local region of the target instance. This local region is called the Region of Competence (RoC). Clustering and distance-based methods were used to define RoC (47,48). The competence of the base models is estimated using samples in the RoC according to predefined criteria.

Several DES algorithms were used in this study:

- K-Nearest Oracles Eliminate (KNORA-E)
- K-Nearest Oracles Union (KNORA-U)
- Dynamic Ensemble Selection-Kullback-Leibler divergence (DES-KL)
- Randomized Reference Classifier (RRC)
- Meta-learning for dynamic ensemble selection (META-DES)

A description of the listed algorithms is given in (49–53). The algorithm type was considered a hyperparameter for optimization. META-DES was eventually selected. An optimized and trained bagging ensemble was used as a pool of DES base models.

##### Stacked generalization

Stacked generalization is a technique that combines multiple models to reduce their biases (54). Various models (level-0 models) make classification or regression predictions, and their outputs form a new set of data (55). The new set of data is used as input for another learning model, (level-1 model), being trained through cross-validation to resolve it. Previously described models were utilized as level-0 models, optimized, and trained individually. Three classic models were considered as level-1 candidates: naive Bayes classifier, logistic regression, and decision tree classifier. Finally, logistic regression was chosen by the hyperparameter optimization procedure.

### Evaluation metrics

Accuracy metric was rejected because it can be misleading in an imbalanced dataset. The comprehensive set of performance metrics derived from ROC analysis was employed instead:

- Area under the receiver operating characteristic curve (AUC)
- Equal error rate (EER)
- F1 score
- Macro F1 score
- True positive rate (sensitivity)
- True negative rate (specificity)
- Positive predictive value (precision)
- Negative predictive value (NPV)

The cut-off point for ROC curves was found by maximizing the Youden’s index:

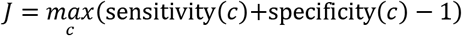

where *J* is the Youden’s index and *c* is the cut-off point.

### Results interpretation

Post-processing of the results was achieved by applying the Shapley additive explanations (SHAP) method. SHAP explains predictions of the arbitrary model using the game theory approach. SHAP computes Shapley values, and features in a dataset are considered as players in a coalition. Shapley value explanation is represented as an additive feature attribution method (56):

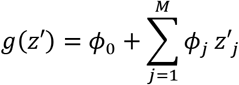

where *g* is the explanation model, *z′* is the coalition vector, *M* is the maximum coalition size, and *ϕ_j_* is the Shapley value for a feature *j*. Using Shapley values, we can obtain an a posteriori feature importance, which is evaluated as an average of the absolute Shapley values per feature (57):

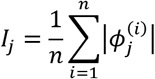

## Results

### Data description

At the feature selection stage, forty-nine significant features were selected out of 827 features, considering the methods described above and the opinions of biomedical experts. The final dataset for model derivation was a table containing 40039 records corresponding to 30550 patients (1108 diabetics and 29442 non-diabetics), with each patient having one or several records in a 5-year timeline. Dataset characteristics corresponding to forty-nine features are presented in Table 1.

**Table 1.**
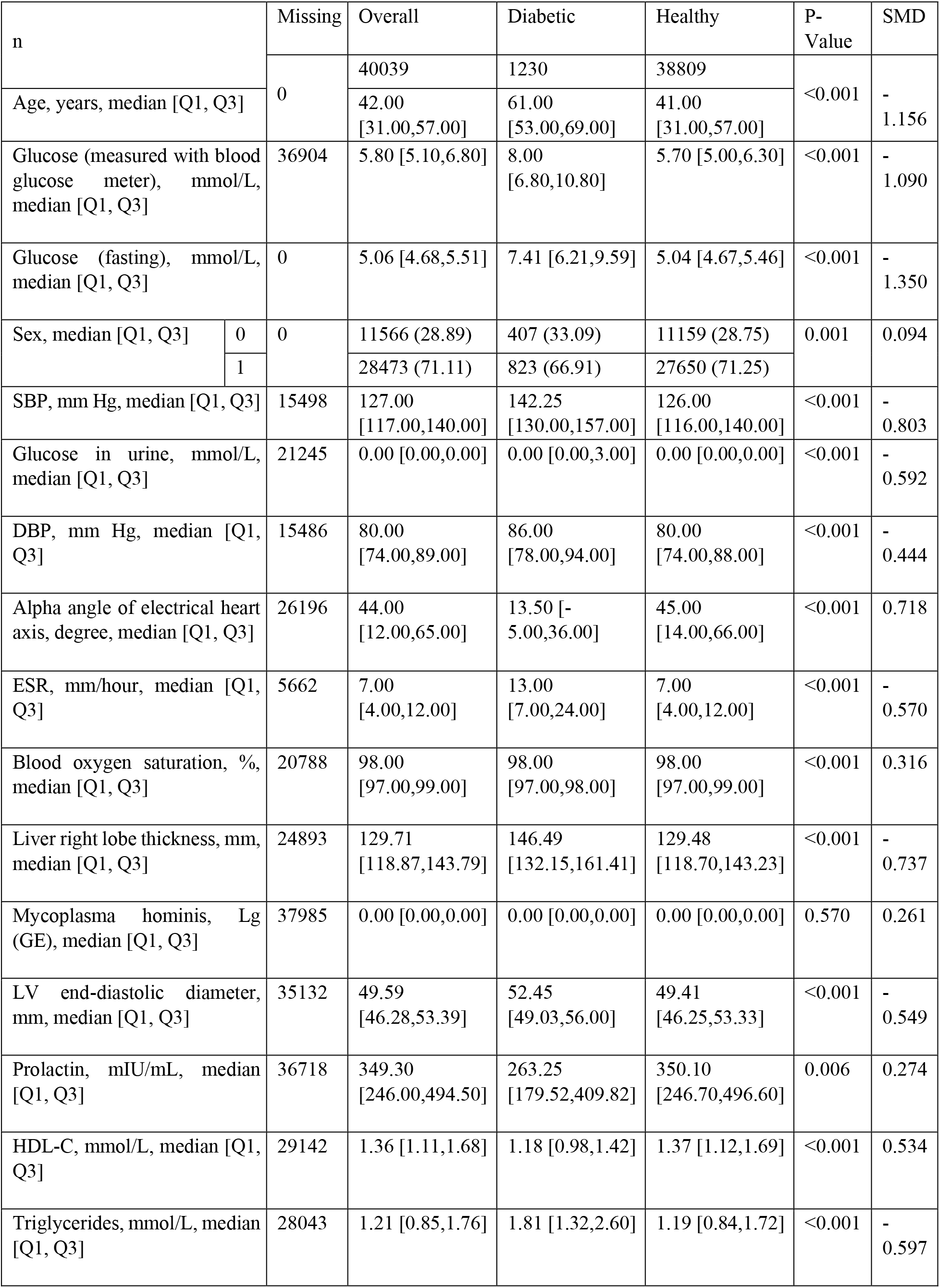
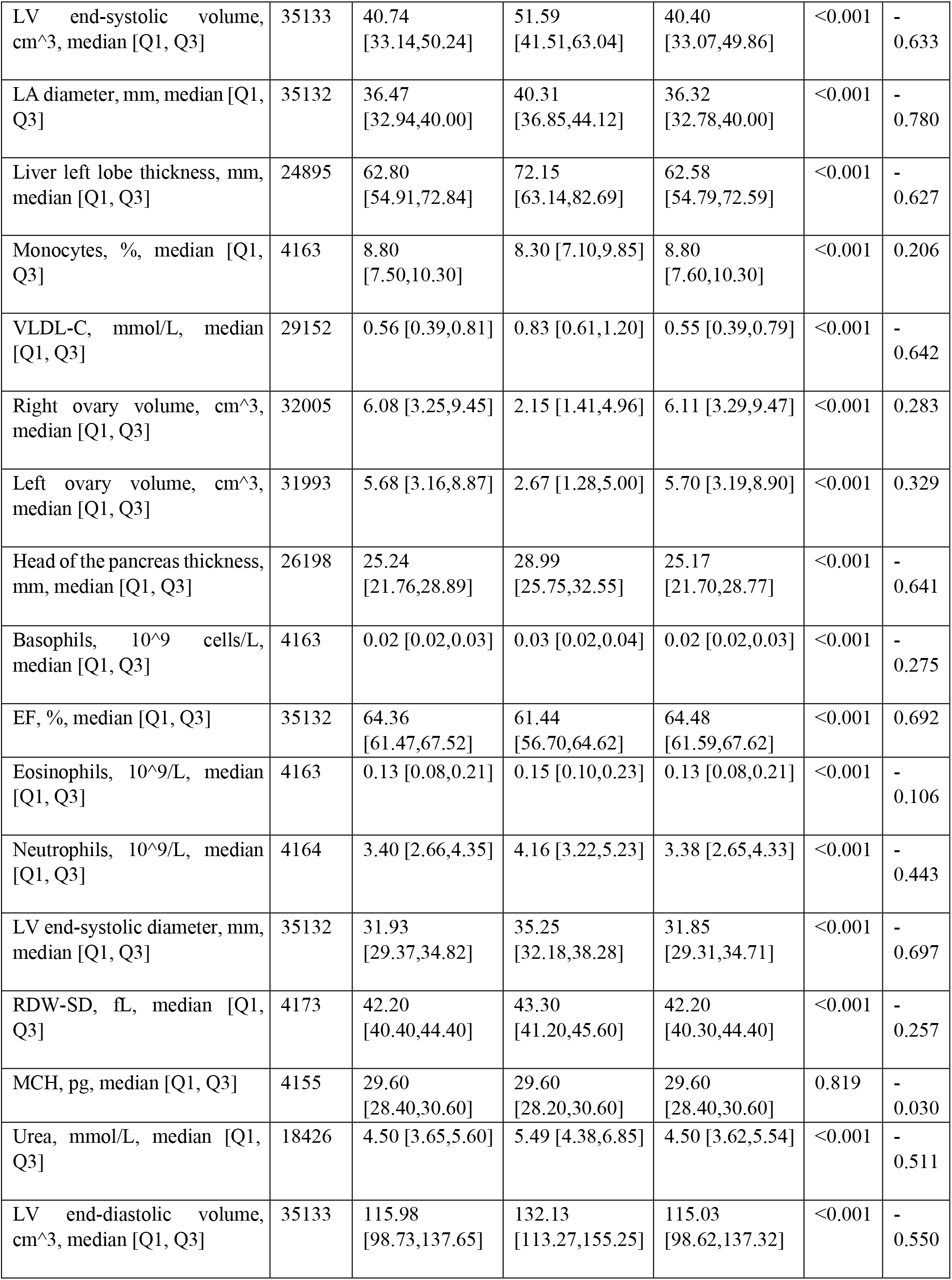
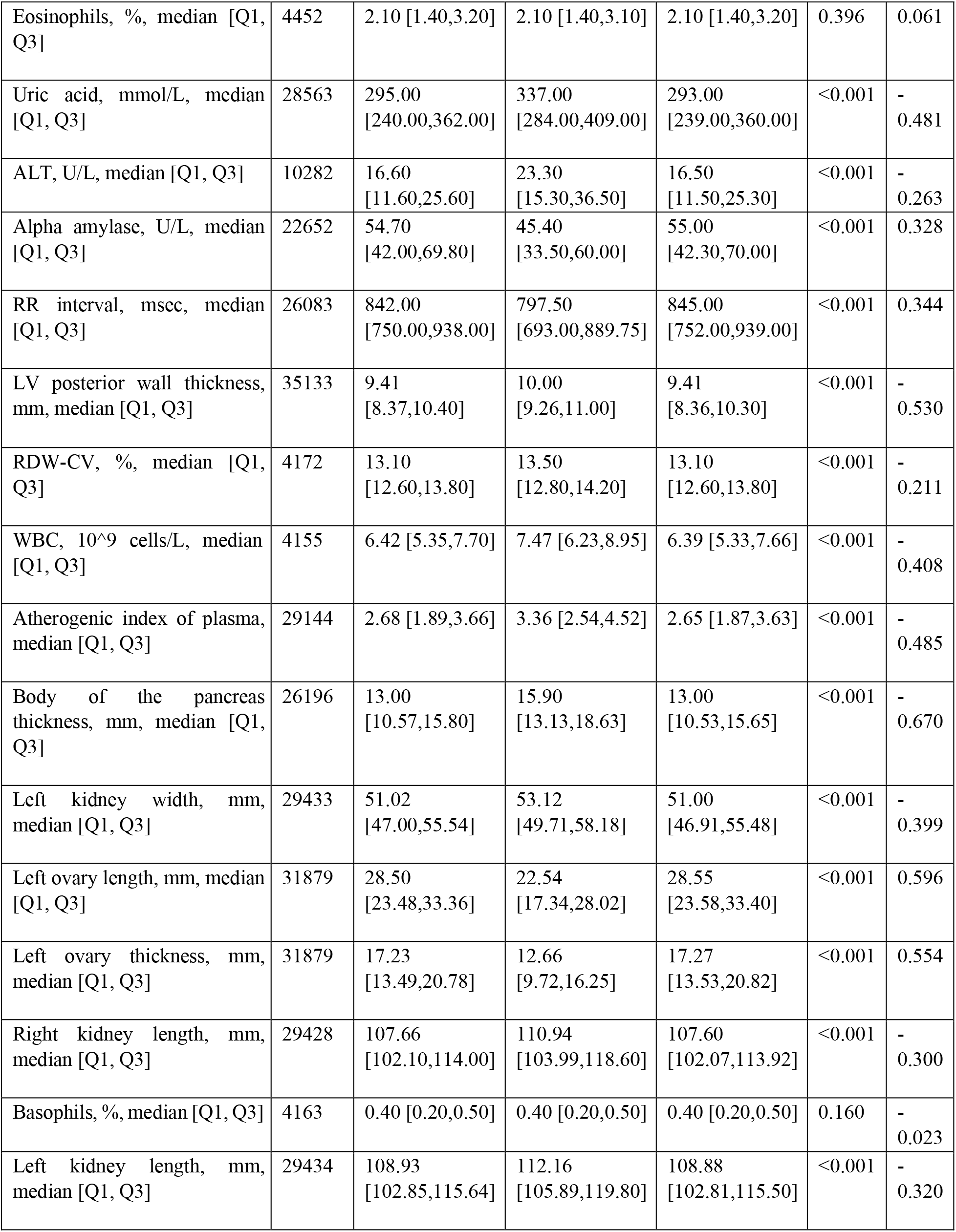
General characteristics of training dataset featuring T2D model predictors. P-values of continuous and categorical variables are measured by the Kruskal-Wallis test and Chi-squared test, respectively. 0 male sex, 1 female sex, SMD standardized mean difference, SBP systolic blood pressure, DBP systolic blood pressure, ESR erythrocyte sedimentation rate, LV left ventricle of the heart, LA left atrium of the heart, HDL-C high-density lipoprotein cholesterol, VLDL-C very low-density lipoprotein cholesterol, EF ejection fraction of the heart, RDW-SD red cell distribution width-standard deviation, RDW-CV red cell distribution width-coefficient of variation, MCH mean cell hemoglobin, ALT alanine aminotransferase, WBC white blood cells.

Individuals with diabetes had greater median age (61 years vs. 41 years for ‘healthy’ individuals), plasma triglyceride levels (1.81 mmol/L vs. 1.19 mmol/L), fasting plasma glucose levels (7.41 mmol/L vs. 5.04 mmol/L). T2D was also associated with hypertension (median systolic blood pressure was 142.25 mm Hg). Moreover, there was a difference in several sonography-derived measurements of the liver and pancreas between the two cohorts.

### Model development and evaluation

The dataset described above was used for evaluating several models. A detailed description of the models is given in Methods (see Machine learning algorithms). Each model was trained and evaluated using training and test subsets, respectively. To obtain more reliable results, we assigned three train/test splits and calculated performance metrics for each of them. The results are shown in Table 2.

**Table 2.** Performance metrics for the first train/test split

|  | Split | AUC | EER | F1 score | Macro F1 score | Sensitivity | Specificity | Precision | NPV |
| --- | --- | --- | --- | --- | --- | --- | --- | --- | --- |
| MLP | #1 | 0.926 | 15.773 | 0.224 | 0.558 | 0.890 | 0.808 | 0.213 | 0.996 |
|  | #2 | 0.960 | 9.716 | 0.346 | 0.644 | 0.915 | 0.893 | 0.213 | 0.997 |
|  | #3 | 0.955 | 10.464 | 0.358 | 0.653 | 0.886 | 0.903 | 0.224 | 0.996 |
|  | Mean | 0.947 | 11.984 | 0.309 | 0.618 | 0.897 | 0.868 | 0.188 | 0.996 |
| Ridge classifier | #1 | 0.946 | 12.836 | 0.430 | 0.698 | 0.789 | 0.940 | 0.295 | 0.993 |
|  | #2 | 0.916 | 16.753 | 0.310 | 0.626 | 0.776 | 0.898 | 0.194 | 0.992 |
|  | #3 | 0.917 | 15.794 | 0.305 | 0.622 | 0.801 | 0.891 | 0.188 | 0.993 |
|  | Mean | 0.926 | 15.128 | 0.348 | 0.649 | 0.789 | 0.910 | 0.226 | 0.993 |
| Ridge classifier with polynomial features | #1 | 0.958 | 10.993 | 0.431 | 0.697 | 0.850 | 0.934 | 0.289 | 0.995 |
|  | #2 | 0.946 | 12.254 | 0.325 | 0.633 | 0.862 | 0.891 | 0.200 | 0.995 |
|  | #3 | 0.947 | 13.001 | 0.256 | 0.583 | 0.902 | 0.837 | 0.149 | 0.996 |
|  | Mean | 0.950 | 12.083 | 0.337 | 0.638 | 0.871 | 0.887 | 0.213 | 0.995 |
| XGBoost | #1 | 0.961 | 10.515 | 0.343 | 0.643 | 0.902 | 0.894 | 0.212 | 0.997 |
|  | #2 | 0.939 | 12.588 | 0.321 | 0.630 | 0.862 | 0.889 | 0.197 | 0.995 |
|  | #3 | 0.934 | 15.121 | 0.363 | 0.658 | 0.809 | 0.916 | 0.234 | 0.993 |
|  | Mean | 0.945 | 12.741 | 0.342 | 0.644 | 0.858 | 0.900 | 0.214 | 0.995 |
| Bootstrap aggregating | #1 | 0.956 | 11.482 | 0.386 | 0.670 | 0.862 | 0.917 | 0.249 | 0.995 |
|  | #2 | 0.933 | 13.484 | 0.272 | 0.596 | 0.874 | 0.856 | 0.161 | 0.995 |
|  | #3 | 0.934 | 14.628 | 0.339 | 0.643 | 0.825 | 0.904 | 0.213 | 0.994 |
|  | Mean | 0.941 | 13.198 | 0.332 | 0.636 | 0.854 | 0.892 | 0.208 | 0.995 |
| Stacked Generalization | #1 | 0.962 | 10.957 | 0.380 | 0.666 | 0.882 | 0.913 | 0.242 | 0.996 |
|  | #2 | 0.955 | 10.925 | 0.374 | 0.662 | 0.878 | 0.911 | 0.237 | 0.996 |
|  | #3 | 0.961 | 10.093 | 0.402 | 0.679 | 0.890 | 0.920 | 0.260 | 0.996 |
|  | Mean | 0.959 | 10.658 | 0.385 | 0.669 | 0.883 | 0.915 | 0.246 | 0.996 |

Application of the DES classifier was declined because it did not provide any metric improvements compared to the base bagging model. ROC curves of the considered classifiers for the first train/test split are shown in Fig. 3. The stacked generalization model yields the best performance results. The stacked generalization model was further improved at the development phase and validated (see below).

**Figure 3.**
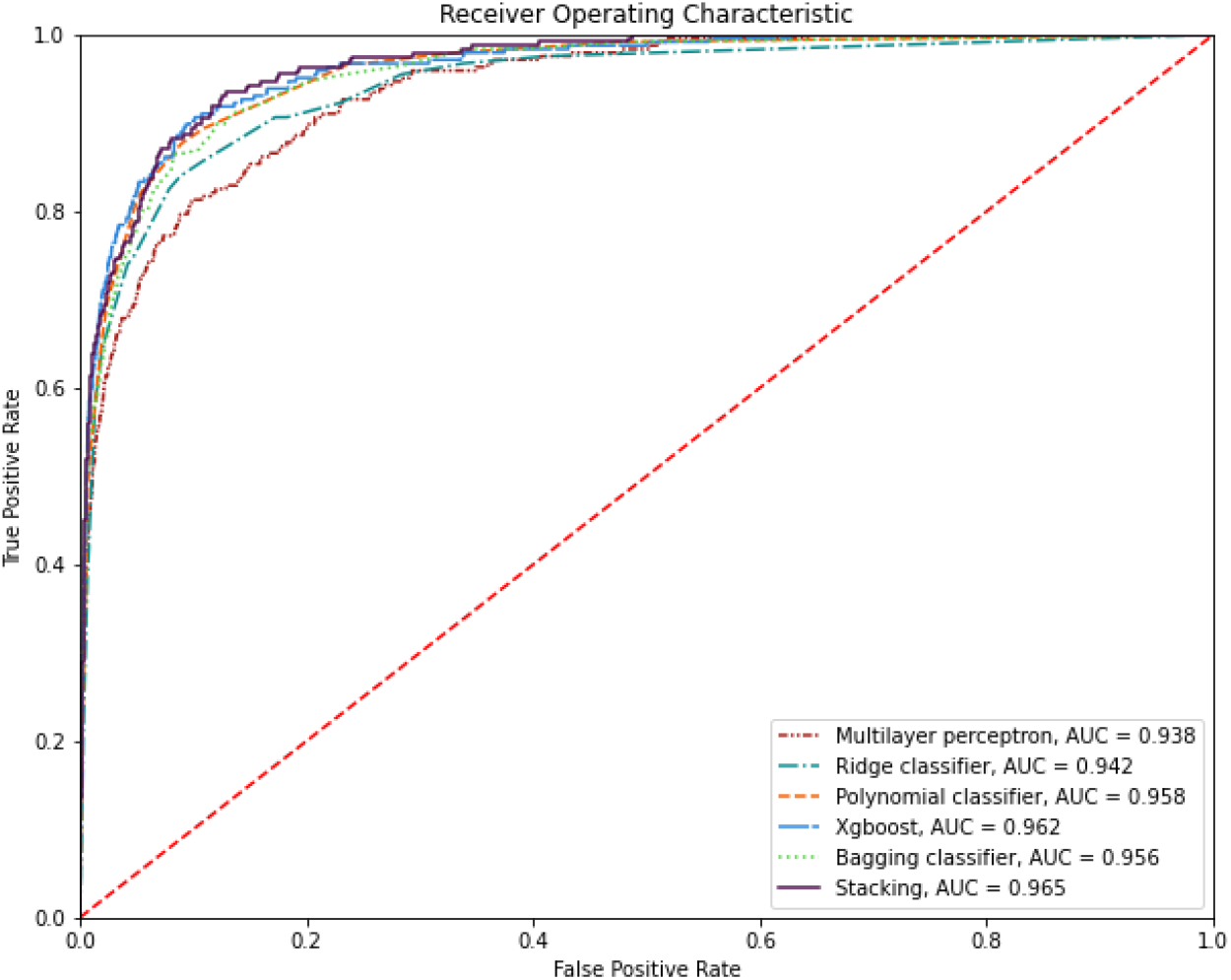
ROC curves for all models

### Interpretation

A post hoc importance analysis has been performed by applying the SHAP method. The beeswarm plot and the bar chart of average absolute SHAP values are shown in Fig. 4A and Fig. 4B. To facilitate interpretation, absolute SHAP values were normalized to add up to one hundred, to simplify interpretation procedures.

**Figure 4.**
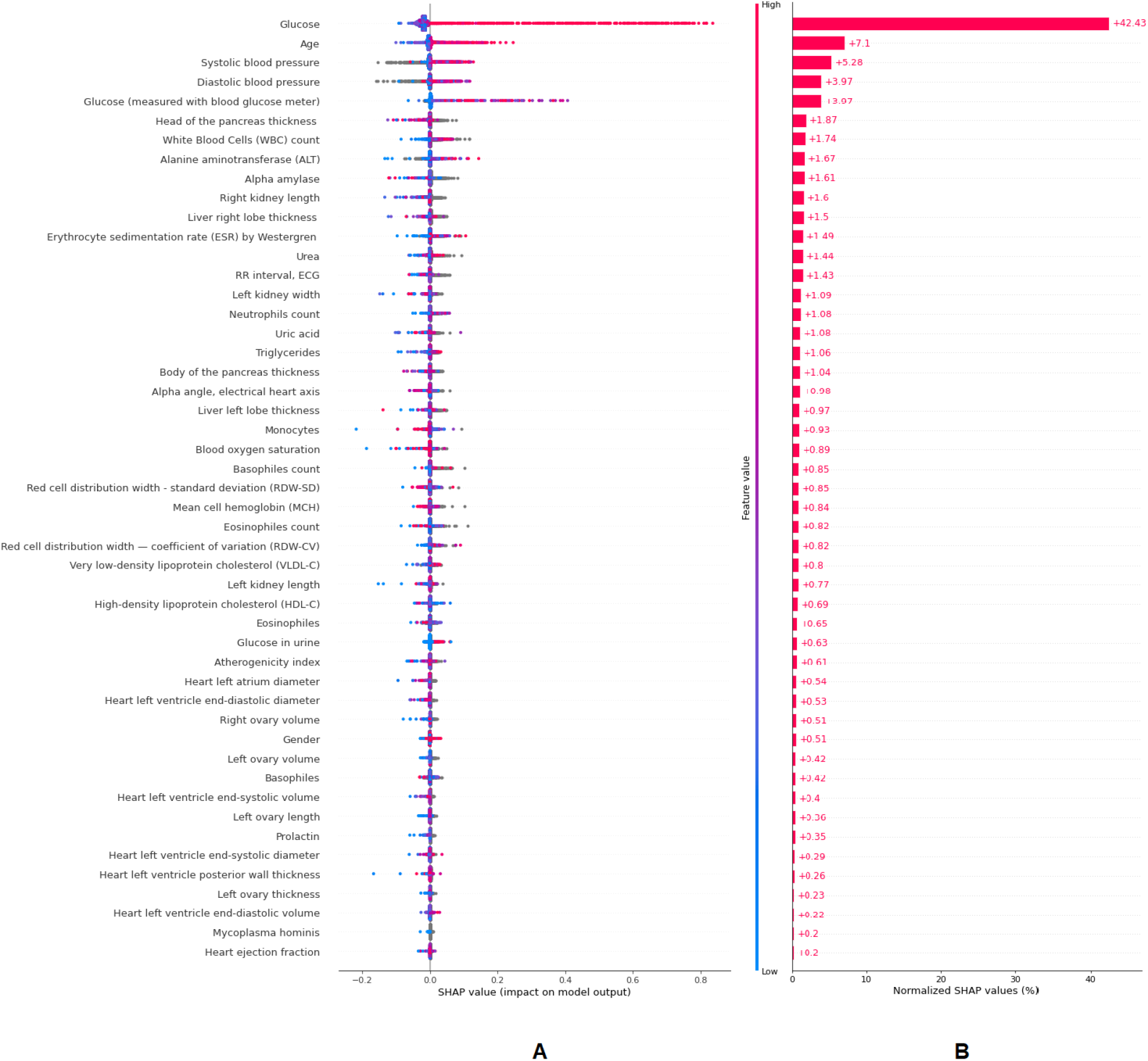
A: beeswarm plots. Each dot corresponds to a patient case. The dot’s position on the horizontal axis shows the impact that feature had on the prediction. B: bar chart of the normalized average absolute SHAP values. Based on these importance values, additional rules were set to employ the model in medical practice. It was proposed to make the five most important features mandatory and to exclude eleven least important ones. This decision makes practical use of the model flexible in case of an insufficient number of non-mandatory predictors (none of them are used in extreme cases), which might be often seen, for example, during initial visits to a clinic, i.e., when EHR system contains a limited number of patient-specific measurements.

### External validation

Data from the National Health and Nutrition Examination Survey (NHANES, USA) was used for external validation of the diabetes model (https://wwwn.cdc.gov/nchs/nhanes/continuousnhanes/default.aspx?Cycle=2017-2020). 2017-March 2020 data considered here represented the data accumulation period for this study. Only features present in the training dataset were selected from NHANES. Similar to the filtration process described above, pregnant women and patients under 18 years old were excluded. The presence of fasting plasma glucose level measurements was mandatory for inclusion in the dataset. The external validation dataset characteristics are shown in Table 3.

**Table 3.**
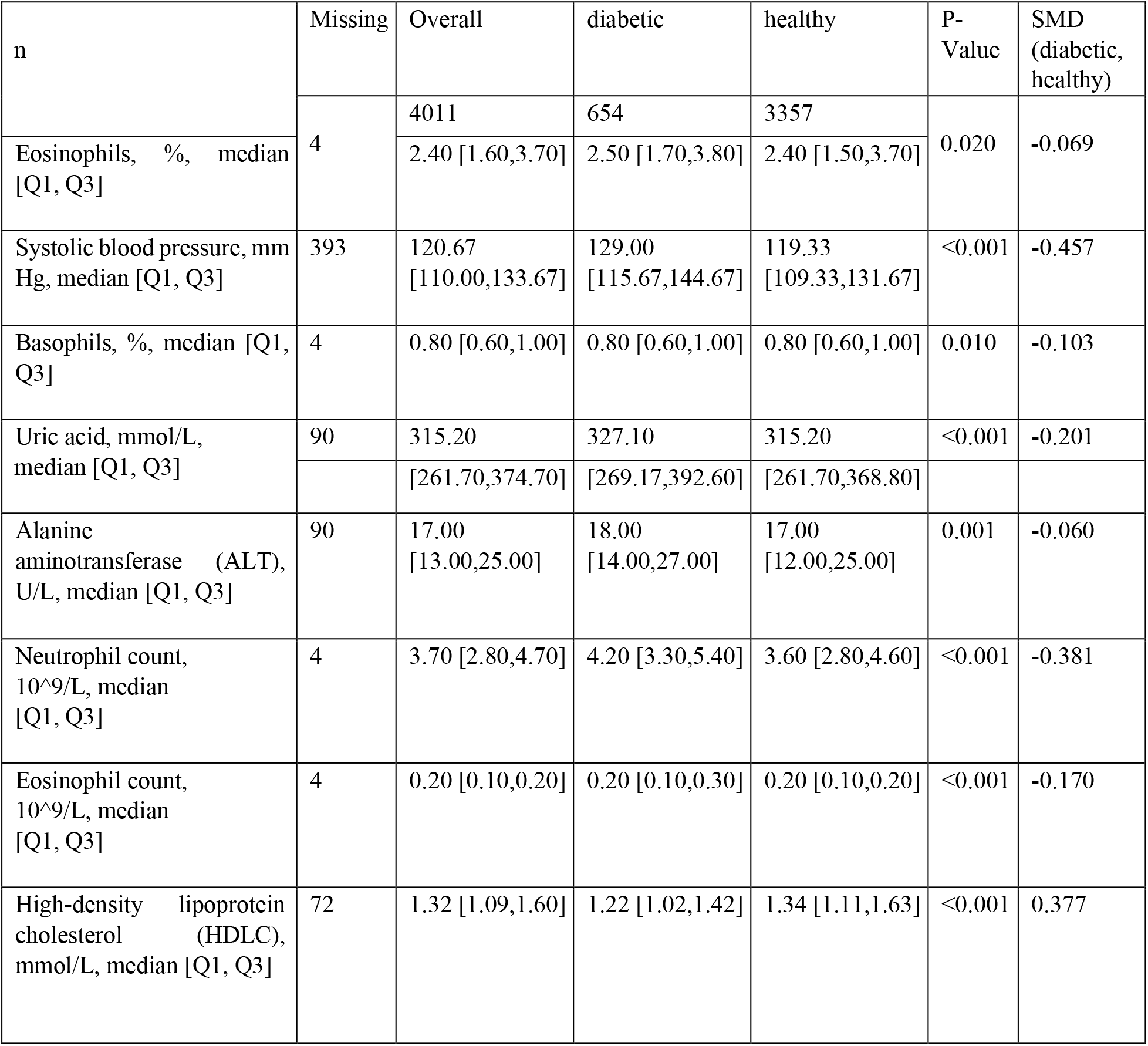
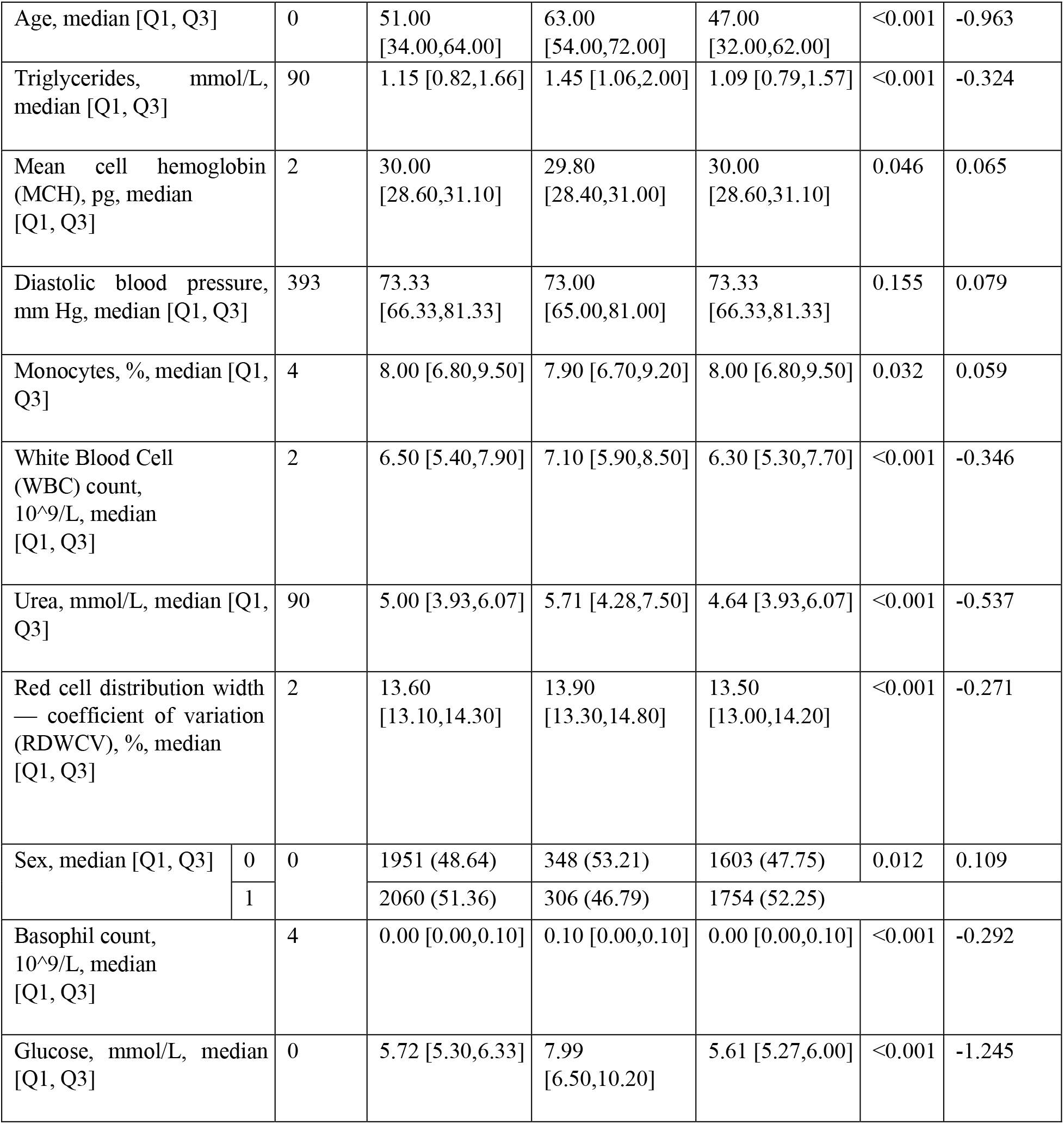
General characteristics of external validation dataset. P-values of continuous and categorical variables are measured by the Kruskal-Wallis test and Chi-squared test, respectively. 0 male sex, 1 female sex, SMD standardized mean difference.

Achieved performance metrics for the external dataset were as follows: 0.899 AUC, 17.896 EER, 0.627 F1 score, 0.764 macro F1 score, 0.809 sensitivity, 0.850 specificity, 0.512 precision, 0.958 NPV. The ROC curve of the stacked generalization model for the NHANES dataset is shown in Fig. 5.

**Figure 5.**
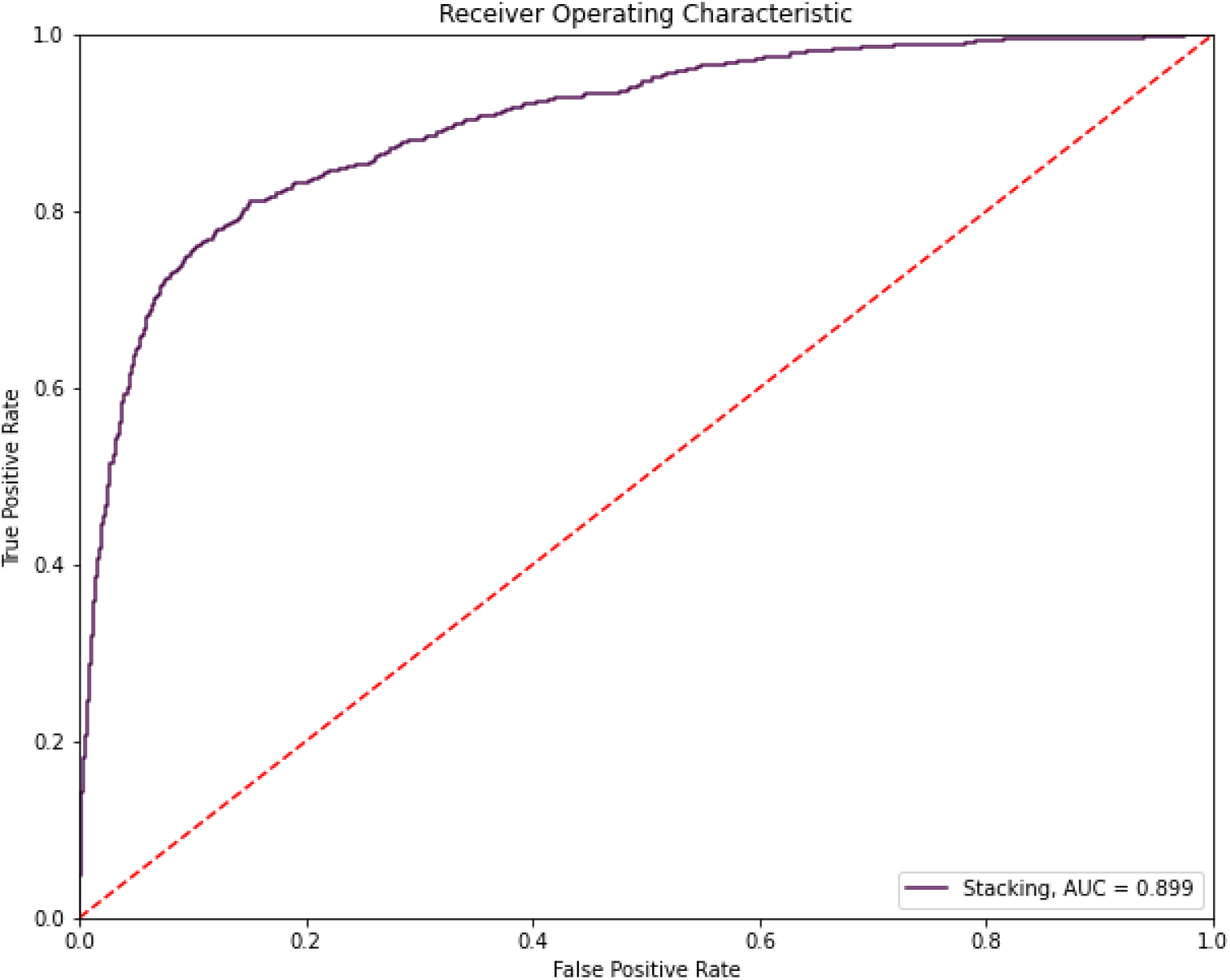
ROC curve for the stacked generalization model (external validation dataset)

## Discussion

Machine learning can be applied to a disease prediction in two separate ways: disease (condition) identification and future occurrence prediction (prognosis). We have developed and validated a model capable of accurate classification of patient-generated data and identification of diabetes-specific patterns in clinical biomarkers values with 0.92 specificity and 0.89 sensitivity (AUC 0.959, precision 0.246, NPV 0.996; see Table 2). Although several clinically recommended diabetes risk scores exist, most of them use virtually the same non-biochemical predictors, such as chronological age, medical history, body mass index. These risk scores demonstrate overall moderate diagnostic accuracy, but are undeniably easy to use in clinical setting, as there is no need to perform most of the laboratory tests (58). One example of such diabetes risk models is the American Diabetes Association diabetes risk score, one of the most well-known clinically validated models. Previously reported performance metrics for the ADA diabetes risk score applied to a NHANES dataset were as follows: 0.79 sensitivity, 0.67 specificity, 0.10 positive predictive value, 0.99 negative predictive value, 0.83 AUC (59). Comparison of NHANES dataset-derived performance metrics for ADA score vs. the model presented in this article proved the latter to be superior in most aspects (see Results: External validation). The difference in the discussed time periods of the NHANES project (1999–2010 period was used in the ADA study) does not invalidate the comparison, considering the uniformity of NHANES data collection and lack of updates on the ADA model performance. Another widely used diabetes prediction model is the Finnish Diabetes Risk Score (FINDRISC). When applied to a European (Spanish) population, the following diagnostic metrics have been reported (based on a fasting plasma glucose criterion): 0.68 AUC, 0.64 sensitivity, 0.64 specificity, 0.076 positive predictive value, 0.976 negative predictive value (60). The ML algorithm proposed in this work can be viewed as a more promising method of detecting undiagnosed diabetes. Due to low positive predictive value of the ADA diabetes risk score and FINDRISC, these models are more suitable for populations with a higher ratio of undiagnosed diabetes, rather than the general population. Trained on general population-derived real-world data and having relatively high positive predictive value, the model presented here is free of this limitation.

Considered here conventional diabetes models rely on non-invasive predictors, such as physical examination information and medical history. These are often supplemented with demographic and questionnaire data. However, laboratory features, being more accurate and unbiased, continue to represent greater interest for models’ development. Recent literature reviews on non-conventional (not clinically used) machine learning tools for diabetes management provided a summary of existing diagnostic ML models that utilize non-invasive features (15,16). Such models demonstrated AUC ranging from 0.83 to 0.90, which is comparable to or higher than that of traditional diabetes models. However, selected best models in the review by Tuppad rely on methods that could be hardly used in clinical settings (e.g., mass spectrometry, ultrasonic sensors) (15). Among the reviewed models, which employ invasively obtained lab features, the highest reported AUC was 0.86. This model based on demographic data (age, sex), biometric data (body mass index, height) and lab features used recurrent neural networks, gated convolutional neural networks, and self-attention networks with transfer learning. The NHANES data for diabetes prediction was used by Dinh et al. who explored a data-driven approach to predict diabetes and cardiovascular disease (61). In that project, laboratory, demographic, dietary, physical examination, and questionnaire information was used to train Logistic Regression, Support Vector Machines, Random Forest Classifier, extreme Gradient Boost, and Weighted Ensemble models. However, contrary to our approach, the features with more than 50% of missing values were dropped out from the final dataset. In diabetes classification, extreme Gradient Boost model achieved an AU-ROC score of 0.957. The study by Dinh et al. and similar works (e.g. (11–13)) do not contain information about external validation, which is crucial to quantify the generalizability of a diagnostic model. In the developed model, fasting plasma glucose level serves as one of the outcome predictors and as an outcome-defining feature (see Methods). There exists a number of studies that avoid doing this because of possible biases (62). However, multiple evidence suggests that glucose and glycated hemoglobin (HbA1c), the two main diagnostic criteria for diabetes, are not as explicit in diagnostic value and need to be interpreted correctly in the context of patient’s medical history, previous glycemic status, complaints, etc. In clinical setting, patients are usually screened for diabetes with a single fasting plasma glucose test, but other conditions, such as acute pancreatitis, may be a cause of elevated glucose levels (63). The phenomenon of stress hyperglycemia, also known as transient hyperglycemia, can be observed in patients without pre-existing diabetes. The association with stress hyperglycemia is common for ischemic stroke, myocardial infarction, and other critical conditions. Sometimes diabetes screening procedure itself may be the cause of stress-induced hyperglycemia (64). Therefore, hyperglycemia at admission does not necessarily correspond to diabetes diagnosis. The baseline data used in this study supports this viewpoint, as shown in Table 4.

**Table 4.**
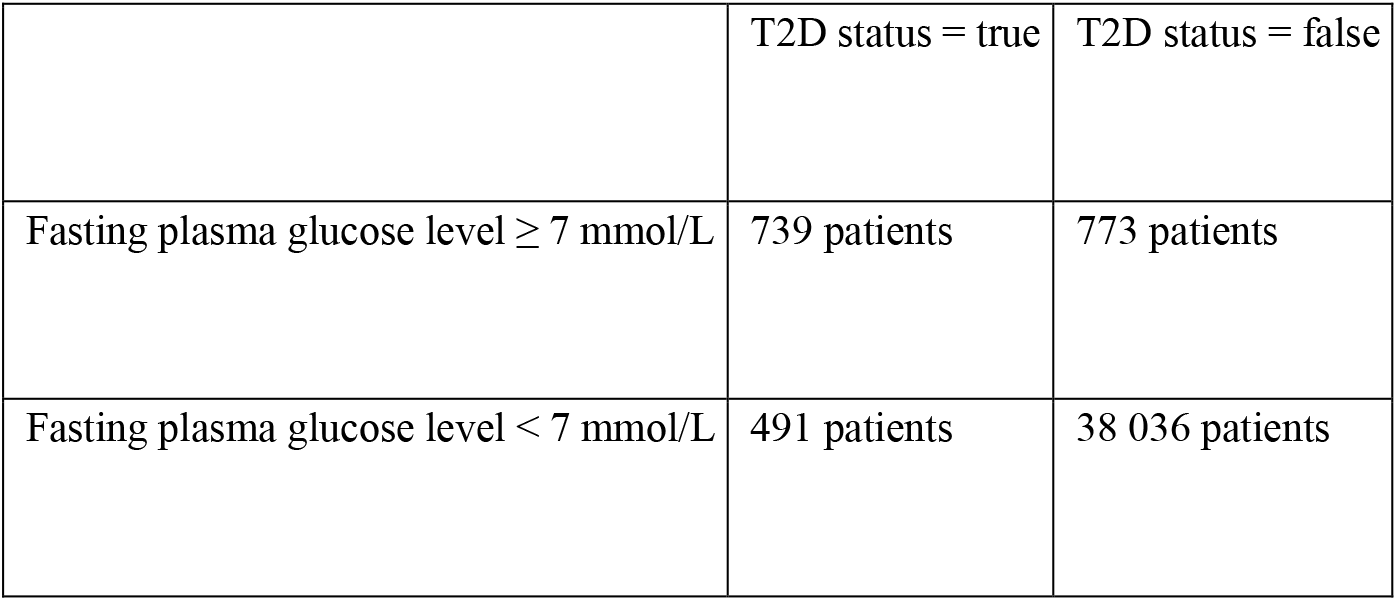
Contingency table of fasting plasma glucose level and diabetes status

|  | T2D status = true | T2D status = false |
| --- | --- | --- |
| Fasting plasma glucose level $\geq 7$ mmol/L | 739 patients | 773 patients |
| Fasting plasma glucose level $< 7$ mmol/L | 491 patients | 38 036 patients |

The odds ratio (OR) calculated from this table equals 74.05 with a 95% confidence interval [64.73, 84.72]. Thus, we confirm the existence of a significant association between high glucose levels and T2D. However, the first row of the contingency table shows that accurate diagnostics of diabetes based solely on fasting glucose levels is not possible.

Similarly, diagnosing diabetes by HbA1c alone may result in many cases remaining undiagnosed because glycated hemoglobin has low sensitivity as a diagnostic biomarker. The sensitivity value of glycated hemoglobin-based diabetes diagnosis was reported to be only 41% (64). Alternatively, the combination of glucose and glycated hemoglobin yields much better results, diagnosing approximately 85% of type 2 diabetes cases (65). Application of two different diagnostic tests to detect diabetes, however, is not obligatory and is recommended by the ADA when diagnosis confirmation is needed.

The ML model, a primary result of this study, can exchange and store data in EHRs, which facilitates its potential practical use in clinical settings. Current ADA guidelines summarize crucial elements of an effective diabetes care framework as the Chronic Care Model (CCM), with decision support being one of the core components of the CCM (see (2) for a detailed description of the CCM). The model presented in this work can serve as a part of the diabetes care framework – a clinical decision support tool in the form of an EHR-based ML-powered screening method for type 2 diabetes in populations of ambulatory patients (symptomatic or non-symptomatic) without confirmed diabetes.

Machine learning approach can be considered as a more appropriate technique for diagnosing and classification of diabetes, since some individuals present with features characteristic to both most prevalent diabetes types at the time of diagnosis, which complicates the classification and prolongs the time needed to choose an appropriate treatment strategy. In view of insufficient understanding of the molecular mechanisms of T2D development, machine learning algorithms are very promising solutions because unknown patterns in data dynamics are recognized and processed. However, ML-based predictions inevitably face the challenge of dealing with imbalanced data during model training. Class imbalance in real-world clinical diabetes data is demonstrated by the incidence rate of diabetes being much lower than that of diabetes-free outcomes., In this study, we did not use the accuracy metric for model performance evaluation because it can be misleading in the imbalanced dataset. A comprehensive set of performance metrics was used instead.

The study has some limitations. First, as misdiagnosis is common in patients with either type 1 or type 2 diabetes, and in view of the absence of genetic data available in electronic health records, there is a minor chance of type 1 diabetes cases to be present in the training dataset (despite the exclusion of ICD codes for T1D). Second, only continuous variables were used as diabetes risk features. Alcohol and tobacco consumption, sleep quality, and other relevant information that requires a questionnaire was not included in the model development. Future versions of the model might consider these risk factors. It should be noted, however, that the definitive contribution of socioeconomic and demographic characteristics of populations to type 2 diabetes risk is yet to be investigated (66).

## Conclusion

We have developed and validated a machine learning classification model for type 2 diabetes that demonstrates several important advantages over established, clinically used methods and possesses a major practical application potential as a part of a clinical decision support framework. To the best of our knowledge, type 2 diabetes ML-model presented in this article is the first diagnostic framework to satisfy all of the following requirements simultaneously: 1) exclusive use of bias-free, objective laboratory medical data; 2) processing of massive arrays of data to identify parameters with the highest diagnostic significance; 3) ability to work with a variable ratio of missing parameters in input data; 4) obtained results that are highly generalizable, having real-world patient-generated medical information as input for the model development.

## Declarations

### Ethics approval and consent to participate

Ethical review and approval are not relevant for this study, because it uses existing data. Informed Consent Statement: not applicable.

### Consent for publication

Not applicable.

### Availability of data and materials

Abbreviated version of the training dataset and processed full external validation dataset are available at https://github.com/biome-science/srep-t2d-data-availability. Web-demo of the diabetes model can be accessed at https://biome-science.com/demo/model/?id=type2diabet. Full data used in model development is available on reasonable request from the corresponding author.

### Authors’ contributions

A.V. and V.G. conceived the experiment, V.D. and V.G. conducted the experiment, V.G. wrote the initial draft, all authors analyzed the results and reviewed the manuscript.

